# How do we respond to the next SARS CoV epidemic/pandemic? A bioinformatics approach with the promise of preventing or reducing the severity of future SARS CoV related pandemics

**DOI:** 10.1101/2024.03.22.586249

**Authors:** Ben Geoffrey A S, Judith Gracia

**Affiliations:** Independent Researcher, Chennai, India; Center of advanced study in Crystallography and Biophysics, University of Madras,Tamil Nadu, India

## Abstract

In this work, we develop a Bayesian weighted scheme to generate evolutionary lineages of a particular viral protein sequence of interest and through a process of clustering and choosing representative lineages from the different clusters according to an evolutionary fitness objective function, we demonstrate it is possible to have anticipated the emergence of the SARS-CoV 2 (2019) strain from the SARS-CoV 1(2004) strain and having shown this retrospectively, we discuss the possibility of applying this approach along with continuous genomic surveillance of SARS-CoVs to prevent or reduce severity of future SARS-CoV related pandemics by being prepared with broad neutralization strategies for anticipated future lineages of SARS-CoVs identified through bioinformatics approaches such as that reported in this work.

## Introduction

The reported work is carried out as a sequel to our previous work [1], wherein we had developed a similar approach of a Bayesian probabilistic weighted scheme to generate mutations more prone and generate mutational lineages of SARS-CoV 2 and evaluate the variants as per an evolutionary fitness objective function and demonstrated that when done so, the computational workflow re-generated the observed micro-evolution of SARS-CoV 2 and thereby establish the feasibility of predicting the micro-evolution of SARS-CoV 2 in-silico based on which therapeutic strategies were discussed for broad protection against the predicted variants. In the presented work, we develop a similar approach of developing a Bayesian weighted scheme to generate the more prone evolutionary lineages of SARS-CoV 1 and among all possible generated evolutionary lineages through a process of clustering the lineages as per similarity among them and selecting representative lineages of each cluster and evaluating them for evolutionary fitness, identify a lineage similar to SARS-CoV 2 in antigenicity and adaptation to infect humans and thereby establish the in-silico feasibility to have foreseen the emergence of a SARS-CoV 2 like strain in antigenicity and adaptation to infect humans from SARS-CoV 1. From the generated evolutionary lineages and their cluster representatives, strains different in antigenicity from that of SARS-CoV 2 but similar in adaption to infect humans were identified and broad neutralization strategies against them are discussed. The scope of using this developed approach in combination with continuous genomic surveillance of SARS-CoVs to foresee and identify future possible lineages of SARS-CoVs with pandemics causing potential and developing broad neutralization strategies against them to prevent or reduce the severity of future SARS-CoV related epidemics/pandemics is discussed. The developed approach is introduced and presented below.

It is understood from the evolutionary history of the Influenza virus, the virus had gained pandemic causing potential by evolutionary reassortments in its antigenic domains and when this novel antigenicity in the virus with sufficient adaptation to infect humans presented itself to an Immunological naïve human population, it resulted in Influenza epidemics/pandemic [2-4]. These learnings from the evolutionary history of Influenza and the understanding of how pandemic causing strains to have emerged in the case of Influenza have been applied to develop a bioinformatic strategy on how to anticipate pandemic causing viral strains in the case of SARS CoVs. The criteria for the emergence of a pandemic causing strain from the evolutionary history of Influenza is understood to be genetic reassortments that allow for the development of novel antigenicity while retaining sufficient adaption to infect humans [2-4]. Mutations are the drivers of the evolutionary lineages of viruses and mutations are of two broad categories which are Single Nucleotide Polymorphisms (SNPs) and Insertions or Deletion (INDELs). By mining UNIPROT data, the most frequently occurring SNPs and INDELs in Spike Glycoprotein of SARS CoVs were identified, and a frequency table was constructed for different types of SNPs and INDELs possible. The frequency table of different possible SNPs and INDELs was used to provide weights to a walker that generates evolutionary emergence for a given protein sequence of interest by generating SNPs and INDELs as per the probabilistic weights from the frequency table of SNPs and INDELs. The Spike Glycoprotein is one of the major antigenic components of SARS-CoVs against which neutralization antibodies are targeted and therefore for the Spike protein of SARS CoV 1(strain associated with 2004 SARS-CoV epidemic), different possible evolutionary lineages were constructed based as per the Bayesian walker that generates mutations as per weights derived out of the UNIPROT data mining.

The generated sequences of the Spike were clustered, and their antigenicity and retention of adaption to infect humans was evaluated. In one of the clusters, a strain was recovered that possessed similar antigenic region/potential of SARS-CoV 2 (2019-nCoV) and ability/adaption to infect humans. The RBD(Receptor Binding Domain) of the Spike Glycoprotein of SARS CoVs are determined to be one of the major antigenic regions targeted by antibodies [6], while also mediating infectivity in humans by being involved in the binding with hACE2(Human Angiotensin-converting enzyme 2) receptor through which cell entry for the SARS-CoVs is mediated in humans [7,8] and the RBD of the generated strain possessed similar affinities for hACE2 and neutralizing antibodies isolated from individuals who were infected with SARS-CoV 2 (2019-nCoV). This demonstrated the ability of the developed approach involving the Bayesian walker which generates different possible evolutionary lineages of the 2004 SARS-CoV 1 strain and when the generated lineages are clustered and shortlisted as per an evolutionary fitness objective of novel antigenicity and adaption to infect humans, it is able to anticipate a strain with similar antigenicity and human adaption/infectivity similar to that of SARS-CoV 2(2019-nCoV). Further, since broad neutralizing is also one of the proposed strategies to mitigate future pandemics [5], other strains generated by the walker which retained adaption to infect humans while having novel antigenicity, different from that of SARS-CoV 2 were identified and for these novel antigenic strains broad neutralization strategies discussed. The results obtained and a future pandemic mitigation strategy thus derived is discussed in the sections that follow along with associated detailed technical methodology.

## Methodology

### Bayesian walker that generates mutation driven evolutionary lineages for a protein sequence of interest

About 14k Spike Glycoprotein sequences of coronaviruses were obtained from UNIPROT and most frequent SNPs and INDELs occurring in them were extracted and a frequency table of most frequent SNPs and INDELs occurring in Spike Glycoproteins of coronaviruses was constructed. The list of UNIPROT IDs of the Spike Glycoprotein Sequences which was used to extract the most frequently occurring SNPs and INDELs in the Spike Glycoproteins of coronaviruses and the extracted frequency table which was subsequently used as weights for the mutation generator is provided in the data link associated with the publication.

A graphical depiction of the frequency plots of representative SNPs and INDELs from the frequency table are show in Fig.1 and Fig. 2 below.

**Fig.1.**
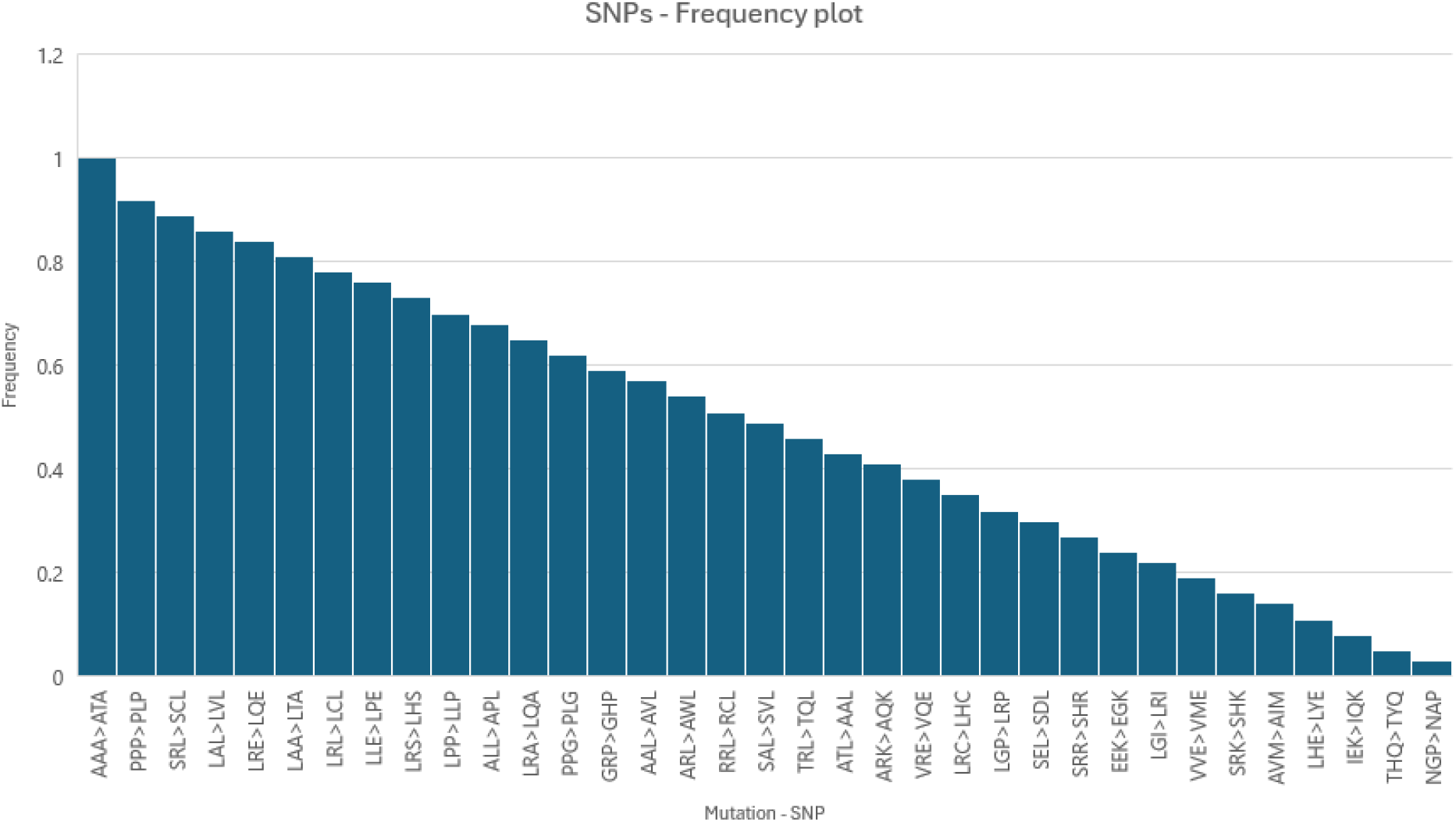
Frequency plot of SNPs

**Fig.2.**
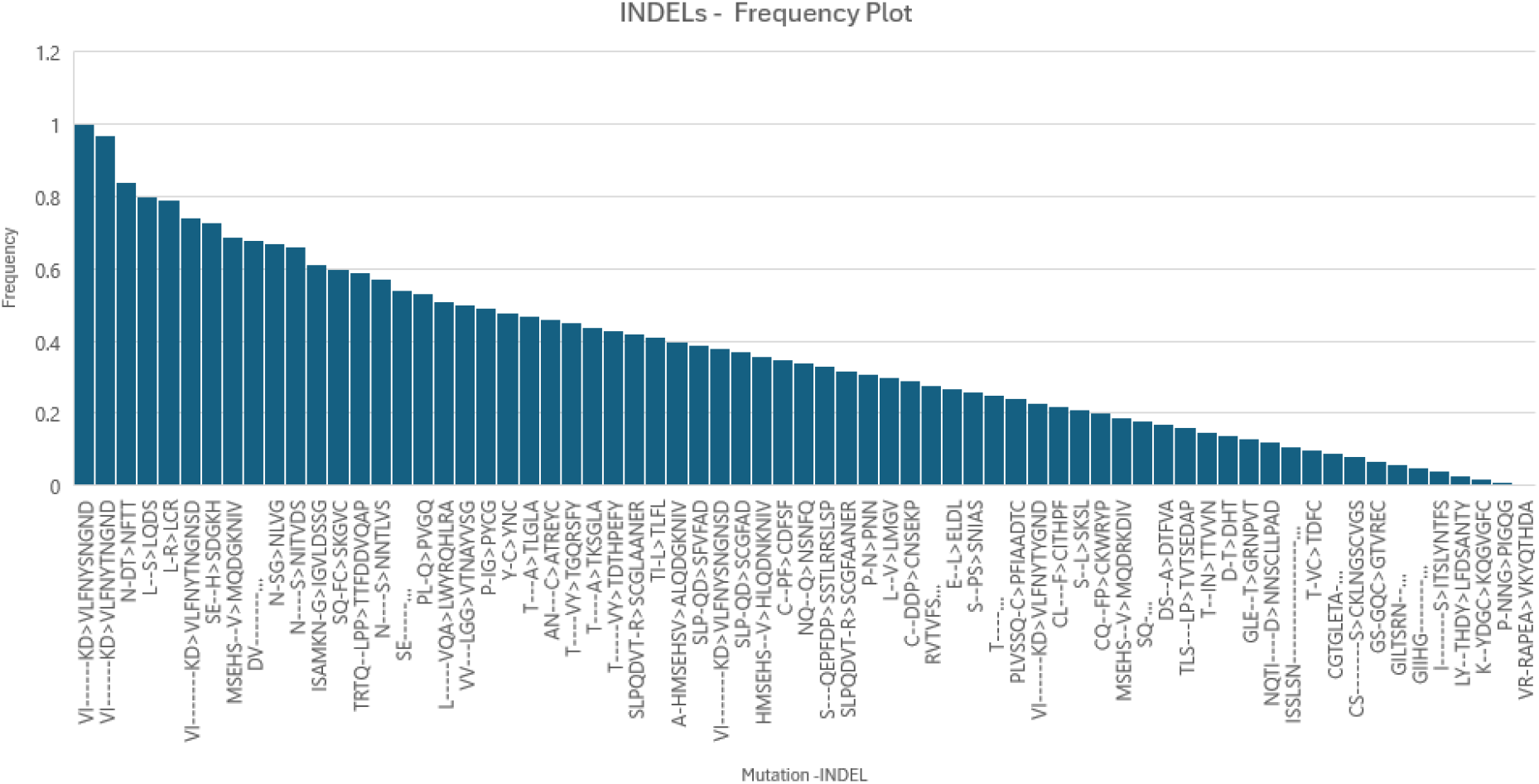
Frequency plot of INDELs

Based on the weights as per the constructed frequency table of SNPs and INDELs observed in the Spike Glycoproteins of Coronaviruses, for the given sequence of interest – Spike protein of SARS CoV 1 (https://www.uniprot.org/uniprotkb/P59594) which is the strain corresponding the 2004 SARS CoV epidemic, its’ evolutionary lineages as per the constructed Bayesian walker was generated to mimic the evolution of SARS-CoV 1 to SARS-CoV 2 in-silico and code used for the generation of sequences as per the Bayesian walker is shared in the Github link associated with the publication.

### Clustering the generated sequence space and identifying cluster representations for evolutionary fitness evaluation

A pairwise sequence similarity matrix was constructed for the generated sequences and the generated sequences were clustered into four clusters as per hierarchical clustering approach and in each cluster, cluster centers were identified which were sequences representative of the cluster. The code used for the clustering is shared in the Github link associated with the publication.

### Molecular modelling workflow that evaluates generated lineages for evolutionary fitness of novel antigenicity through reduced affinity to known neutralizing antibody and retention of adaption to infect humans through affinity for hACE2 binding

The generated chimeric Spike sequences were screened for novel antigenicity and retention of adaptation to infect humans. As discussed earlier, the RBD of the Spike is one of the major antigenic regions recognized by antibodies [6] and the RBD also mediates SARS-CoVs human infectivity through its role in binding with hACE2 for human cell entry of the virus [7,8]. The structure a neutralizing antibody isolated from SARS-CoV 2 infected patients bound to the RBD of SARS-CoV 2 was identified from RCSB-PDB with PDB ID : <u>7CDI</u> and used as a template in homology modelling carried out in MODELLER[9], to model the structure of RBDs corresponding to the chimeric Spike sequences generated from the approach and understand the novel antigenicity in the RBDs of the generated Spike sequences through their reduced affinity for presently known neutralizing antibodies and their Immune escape therein and increased potential to affect an Immunological naïve population through their novel antigenicity. To quantify the decrease in affinity of the RBDs corresponding to generated Spike sequences for the neutralizing antibodies, the RBD-antibody complexes were analyzed through the tool SURFACE [10] which provides the interaction energy between the RBD and the neutralizing antibody for the given RBD-antibody complex. Similarly, to model the retention of adaption in the generated Spike sequences to infect humans through human cell entry mediated through RBD-hACE2 interaction, the RBD-hACE2 complex was identified from RCSB PDB with PDB ID : <u>6M0J</u> and used as a template to model RBD-hACE2 complex corresponding to the generated Spike sequences and retention of the affinity of RBD for hACE2 in the generated Spike sequences was assessed through interaction energy scores obtained for the RBD-hACE2 complexes from the tool SURFACE[10]. In the developed approach, Molecular dynamics simulation is also as technique to assess the stability of the protein structure corresponding to the generated Spike sequences. GROMACS [11] bio-molecular simulation package was used to perform a classical molecular dynamics simulation of the 3D structure corresponding to the generated Spike sequences. The AMBER [12], amber99sb-ildn was force field of choice for the protein and the system was solvated in a solvent box with ions added to physiological pH and to mimic physiological temperature and pressure heated in NVT and NPT ensembles to 300K and 1 bar pressure respectively in a 100 ps NVT and NPT runs with the temperature and pressure controlled by Berendsen thermostat and barostat respectively [13]. This was followed by 50 ns production run in which the stability of the protein was assessed by observing the stabilization of the protein RMSD in time. The stable part of the trajectory was chosen for MMPBSA based binding energy calculation for the RBD-antibody and RBD-hACE2 complexes carried out using the gmx_mmpbsa tool [14]

Github code link reference in the methodology : https://github.com/bengeof/SARS_CoV_Evolution_Generator

Data link referenced in the methodology : https://drive.google.com/drive/folders/1lkPHixAqDX43UBjox_VVaQ_LedYR8KwB?usp=sharing

The results obtained are presented and discussed in the section that follows.

## Results and Discussion

The generated chimeric Spike sequences belonging to the 4 different clusters is shared through the data link associated with this work. From among the clusters derived from the generated sequences, a particular strain was identified which had similar antigenicity to that of SARS-CoV 2 in that, the RBD neutralizing antibody of SARS-CoV 2, P2C-1F11[15], showed comparable affinity for RBD of the generated sequence and the RBD of the generated sequence had similar affinity for hACE2 as that of SARS-CoV 2. This provides validation to the approach that from the generated evolutionary lineages of SARS-CoV 1 as per the Bayesian walker, it is possible to identify a strain similar to SARS-CoV 2 among the sequences shortlisted as per the approach developed in this work. Further, Spike sequences with novel antigenicity, different from that of SARS-CoV 2 were also identified from the approach. The overall protein structure stability corresponding to these generated sequences was evaluated using classical molecular dynamics and the binding energy profiles of the RBD-hACE2 and RBD-antibody(P2C-1F11) complexes were calculated to evaluate their retention of ability to infect human cells through hACE2 binding and Immune evasion through reduced affinity for known antibodies. The generated Spike sequences belonging to the four clusters can be accessed at the data link shared below. Spike Sequence identified from these four clusters with similarity to the antigenicity and human infectability as that of SARS-CoV 2 has been named Spike1 and it’s corresponding Receptor Binding Domain as RBD1, while the 3 other Spike sequences identified from the 3 other clusters with novel antigenicity, different from that of SARS-CoV 2 and retention of ability to infect humans by possessing similar affinity for hACE2 as that of SARS-CoV 2 have been name Spike2, Spike3, Spike4 and their corresponding Receptor Binding Domains as RBD2, RBD3 and RBD4 respectively.

Data link referenced in the results section: https://drive.google.com/drive/folders/1WySujDxECZQ6p3KrN6XNcimPdCoFByMt?usp=sharing

The stability of the Protein structure associated with Spike1, Spike 2, Spike 3 and Spike 4 sequences was evaluated through a 50-nanosecond classical molecular dynamics simulation and their stability indicated by means of the protein RMSD stabilization is graphically depicted in Fig.3 below

**Fig 3.**
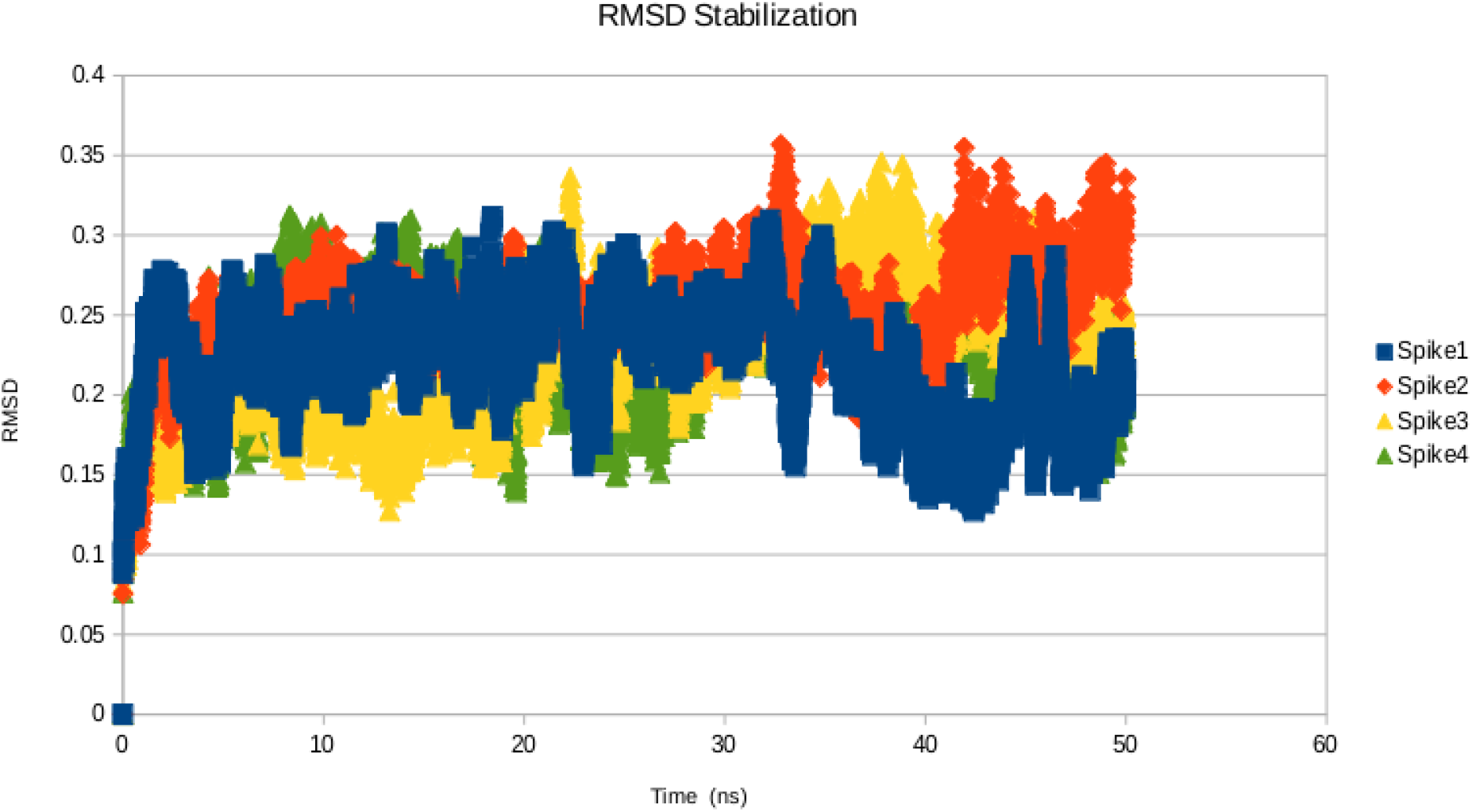
RMSD stabilization indicating the stability of the structure corresponding to the chimeric Spike sequences generated through the approach.

The SURFACE tool-based interaction energy and MMPBSA based binding energies of the RBD-hACE2 and RBD-antibody are tabulated in Table 1 and Table 2 below respectively and the RMSD stabilization of the RBD-hACE2 and RBD-antibody complexes corresponding to Spike1, Spike2, Spike3 and Spike4 is shown below in Fig.4 and Fig.5 respectively.

**Fig.4.**
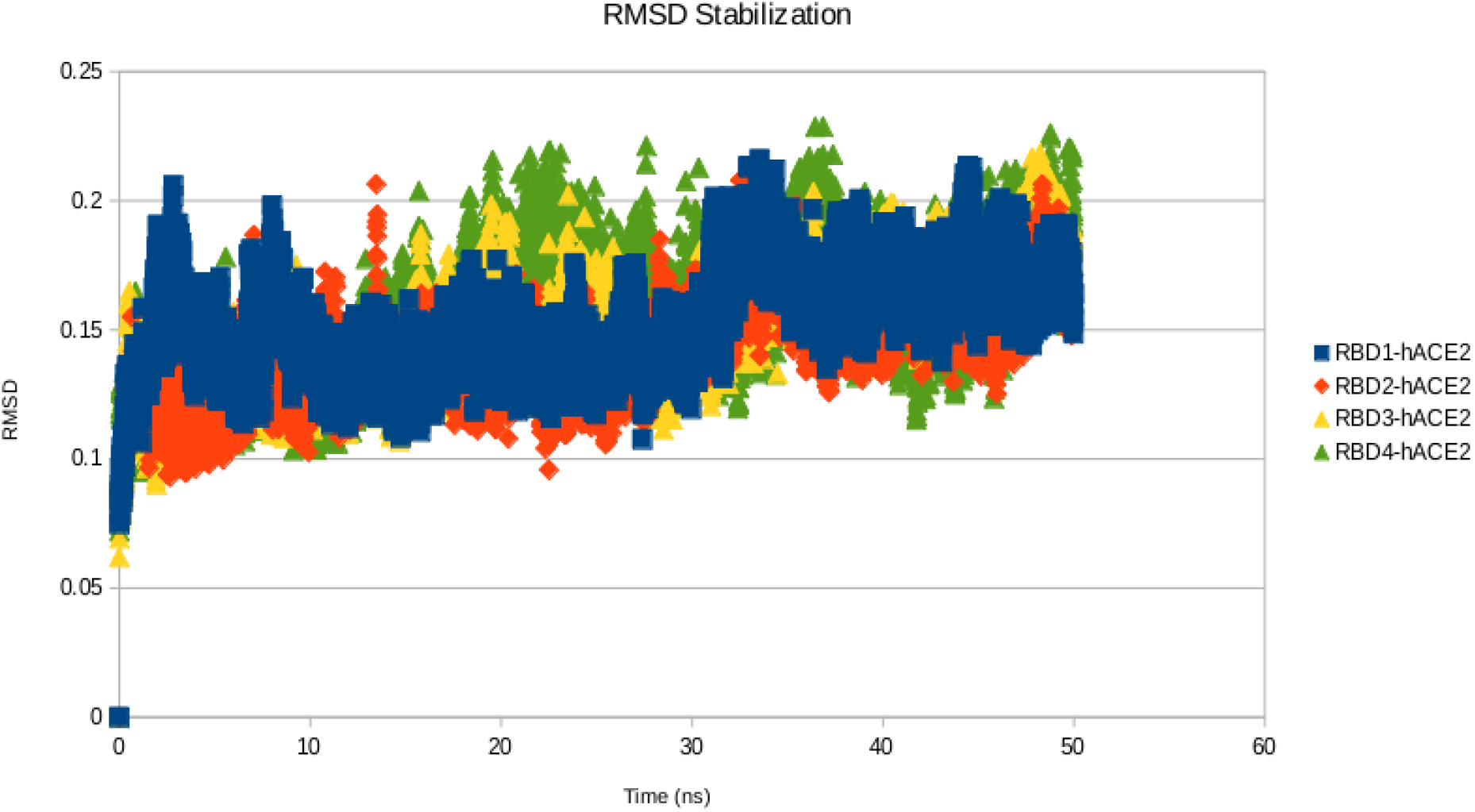
RMSD stabilization of RBD-hACE2 complexes corresponding to the Spike sequences generated by the Bayesian walker and shortlisted through the developed approach

**Table 1.**
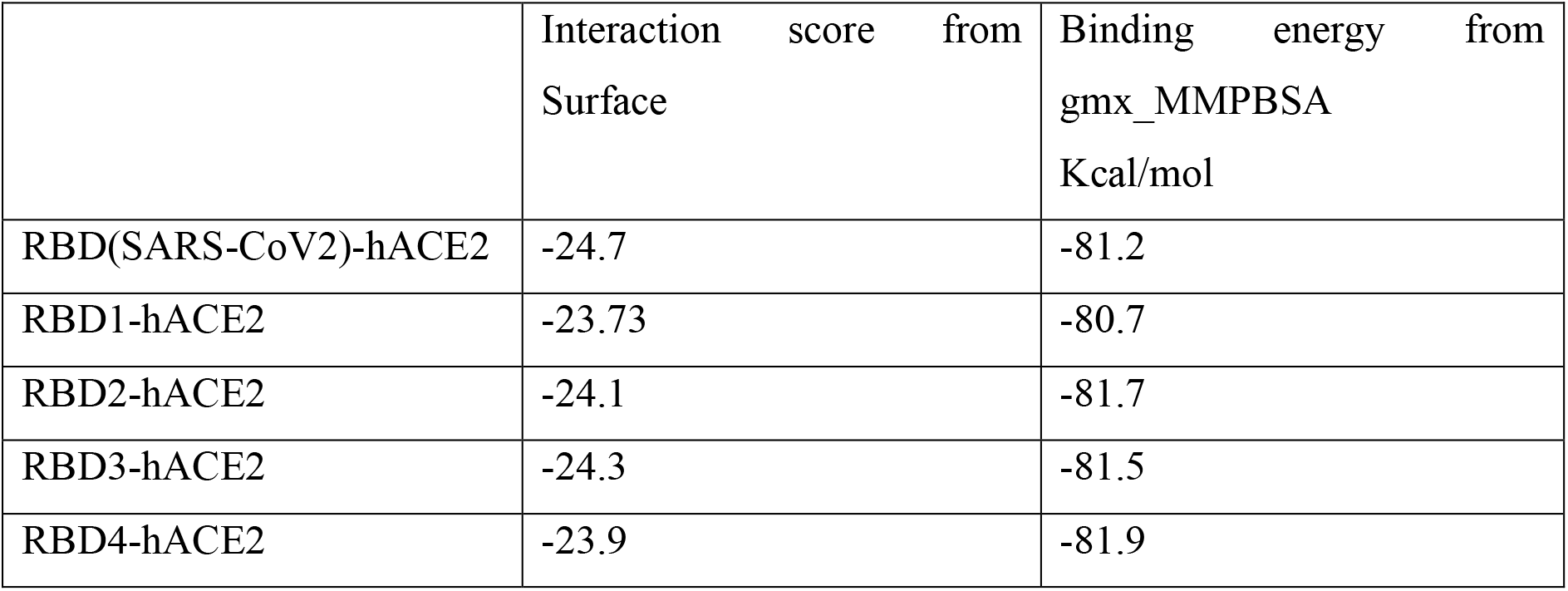
Interaction and Binding energies of RBD-hACE2 complexes.

**Table 2.**
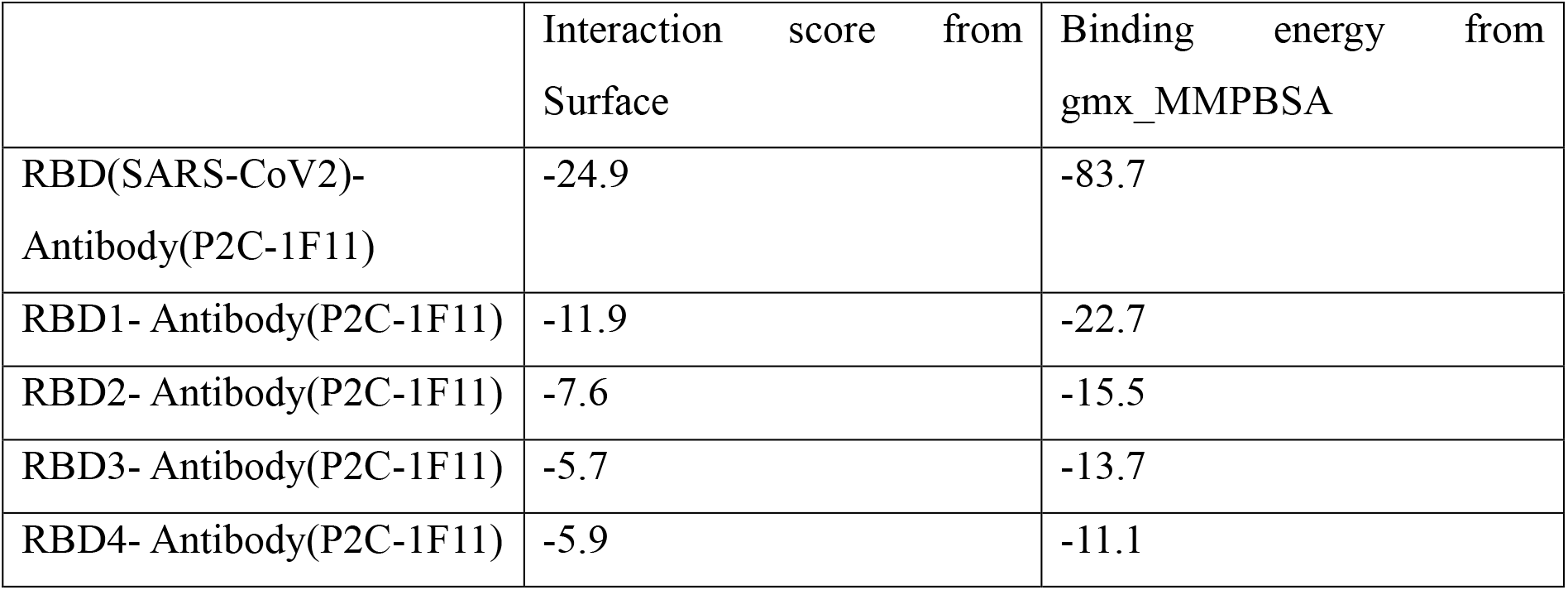
Interaction and Binding energies of RBD-Antibody(P2C-1F11) complexes.

The data indicates the RBD1 corresponding to Spike1 has similar affinity for hACE2 and RBD neutralizing antibody P2C-1F11 as that of SARS-CoV 2. This indicates the ability of the approach to have generated and identified a strain similar to that of SARS-CoV 2 among the possible evolutionary lineages generated for SARS-CoV 1 by the Bayesian walker approach. Further, when all the possible evolutionary lineages of SARS-CoV 1 as per the Bayesian walker were generated and clustered and representative sequences were selected from the clusters, Spike sequences with novel antigenicity inferred through their reduced affinity for SARS-CoV 2 neutralizing antibody P2C-1F11 and retention of adaption to infect humans through retention of affinity for hACE2 were identified. An approach such as a cocktail of monoclonal antibodies prepared against the Spike Sequences namely Spike1, Spike2, Spike3 and Spike4 identified by the approach among the generated evolutionary lineages of SARS-CoV 1 could have prevented or reduced the severity of the pandemic seen in 2019 by having a cocktail of monoclonal antibodies with broad neutralization ability including ability to neutralize the strain corresponding to the 2019 SARS-CoV 2 with RBD antigenic region of Spike1 being similar to SARS-CoV 2, while also having broad neutralization ability. The alignment of the RBD regions of the chimeric Spikes generated and shortlisted in this approach of Spike1, Spike2, Spike3 and Spike4 bearing the names RBD1, RBD2, RBD3 and RBD4 are compared with RBDs SARS-CoV 2(2019 strain) and SAR-CoV 1(2004 strain) and their Clustal Omega [16] alignments obtained can be accessed in the data link associated with the publication and is shown in Fig.6 below

**Fig.5.**
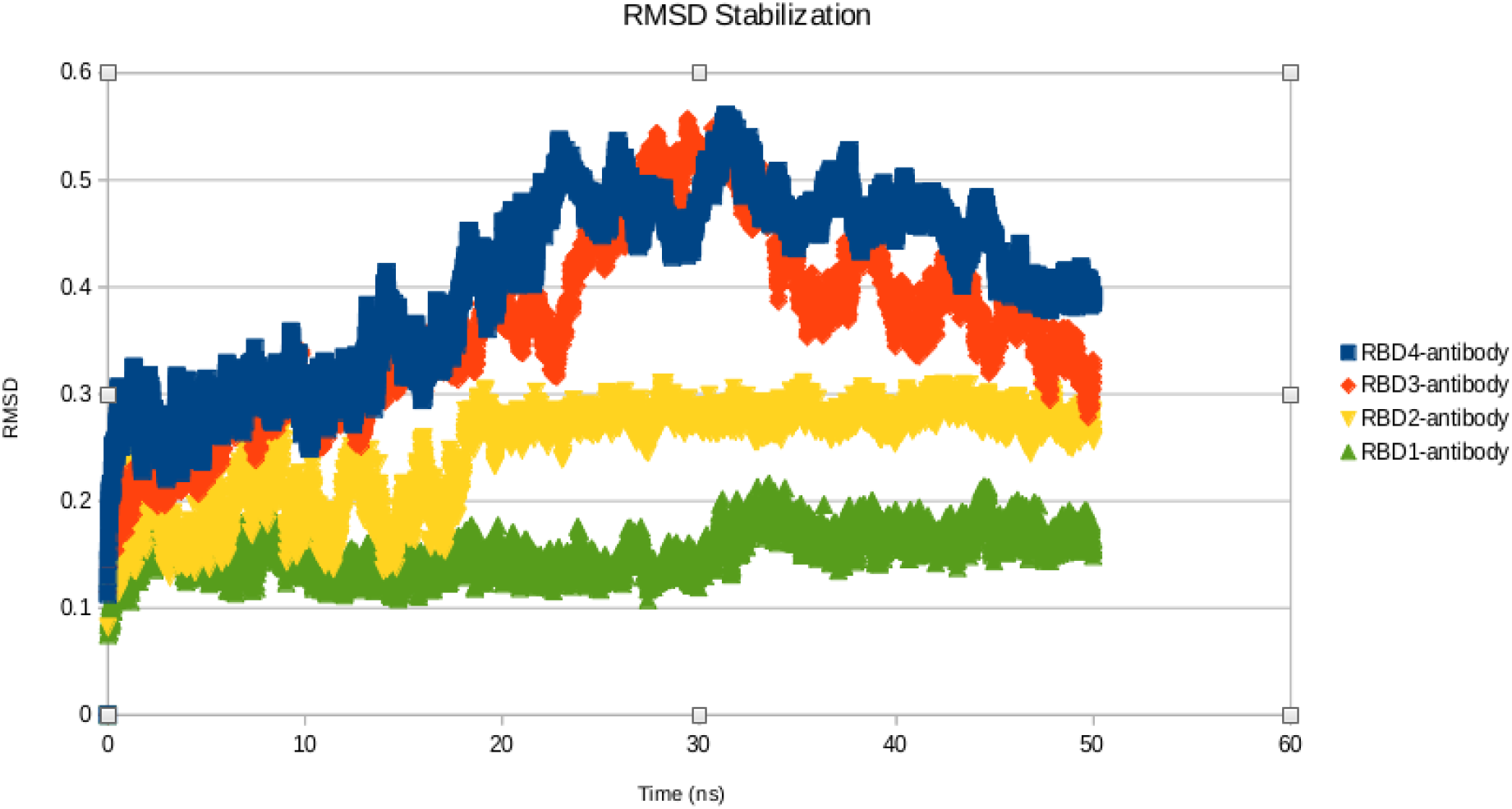
RMSD stabilization of RBD-antibody complexes corresponding to the Spike sequences generated by the Bayesian walker and shortlisted through the developed approach

**Fig.6.**
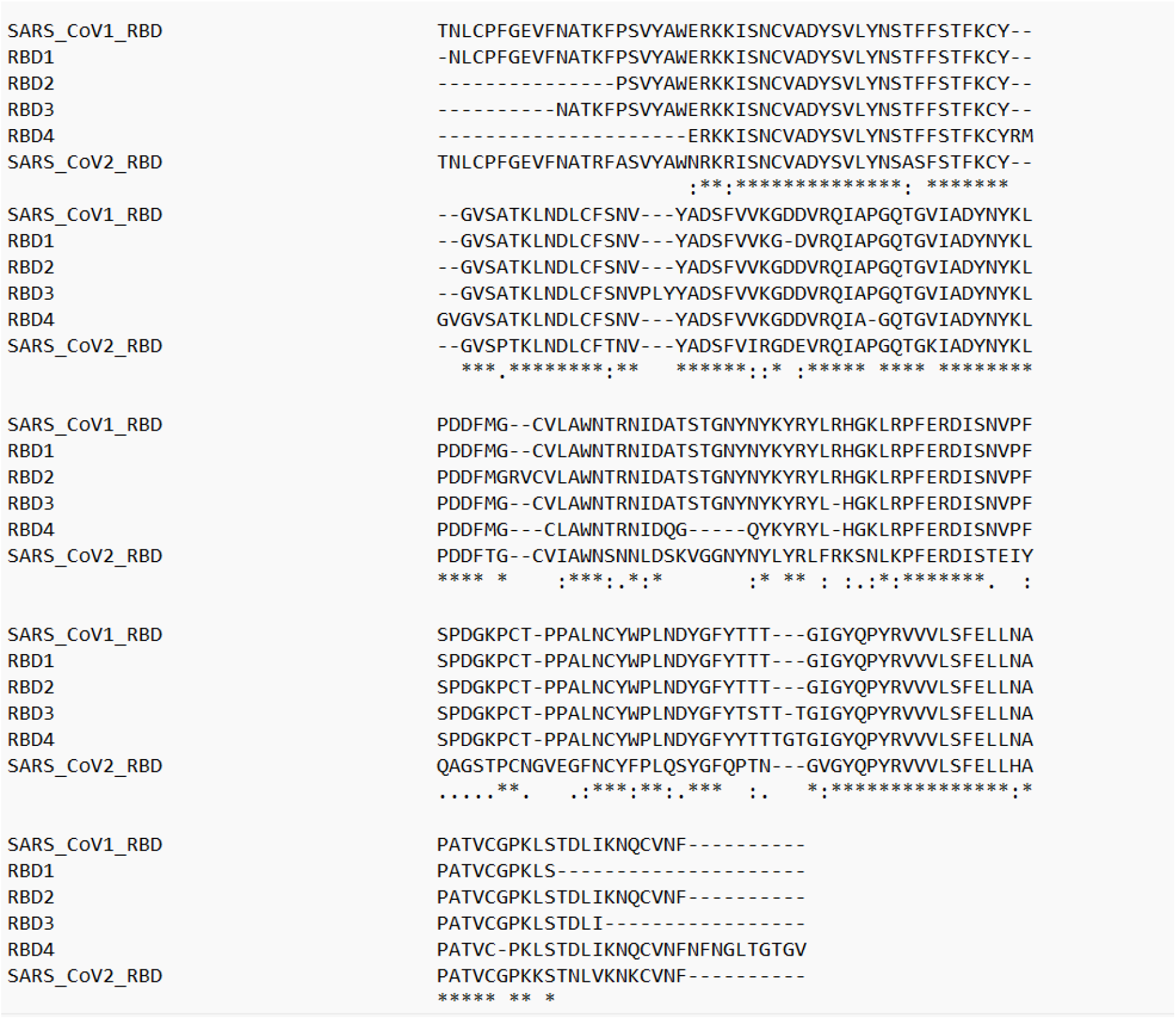
Alignment of RBDs

Thus, a combination of continuous genomic surveillance of CoVs and an approach such as that developed in this work of generating future possible evolutionary lineages for dominant SARS-CoV lineage as of today and shortlisting the future possible evolutionary lineages through clustering them via similarity and among diverse clusters, selecting cluster representative sequences that satisfy the criteria of evolutionary fitness of novel antigenicity and retention of human infectability, and by having a broad neutralizing strategy developed against such future possible lineages, it will be possible to prevent or reduce the severity of future SARS-CoV related pandemics/epidemics. The scope of extending this approach beyond application to SARS-CoVs is also discussed below.

By studying the origins of previous pandemics, it is understood the viral pandemics have emerged when the virus undergoes sufficient genetic re-assortments such that it gains novel antigenicity while maintain human infectability posing a pandemic threat to population which is Immunological naïve to the antigenic threat posed by the virus. Further, viruses infecting animals undergoing continuous genetic re-assortments can gain ability to jump the species barrier and infect humans. Therefore, a combined strategy of continuous genomic surveillance of strains of viruses circulating in animals and humans to track the on-going genetic re-assortments and having neutralization strategies for future possible lineages that have epidemic/pandemic causing potential foreseen by computational means by tools such as that developed in this work will help in prevention or reduction of severity of such future epidemic or pandemic occurrences.

## Conclusion

By mining UNIPROT data, the mutations and genetic re-assortments which occur with more frequency in the Spike protein of Coronaviruses were identified and based on which a Bayesian generation scheme was created which generates evolutionary lineages through mutations which are more prone for Spike protein sequence of interest by generating mutations as per the frequency table of mutations derived from the data mining. We demonstrate that when the approach of generating evolutionary lineages developed in this work is applied to generate evolutionary lineages of the Spike protein of SARS-CoV 1(2004 strain) and when the generated evolutionary lineages are clustered and when representative sequences of the clusters are chosen and evaluated for evolutionary fitness, it was possible to identify a strain similar in antigenicity and adaption to infect humans as that of SARS-CoV 2(2019 pandemic strain) while also identifying novel chimeric Spike Sequences which had diverse antigenicity differing from SARS-CoV 2 and each other while retaining the ability to infect humans. While this was carried out retrospectively, we discuss a forward-looking aspect of this work wherein for SARS-CoVs lineages dominant as per genomic surveillance of today, the developed approach can be applied to foresee the emergence of SARS-CoV lineages with future epidemic/pandemic threat and by developing broad neutralization strategy against these anticipate lineages with epidemic/pandemic threat, the severity of such future SARS-CoV related epidemics/pandemics can be reduced.

## Supporting information

Images and Tables

## References

1. Ben Geoffrey, A. S., and Judith Gracia. “A Bayesian walker coupled with a computational workflow that generates the micro-evolution of SARS-CoV-2 and makes predictions of new mutations that can emerge.” Journal of Biomolecular Structure and Dynamics (2023): 1–9.

2. Worobey, Michael, Guan-Zhu Han, and Andrew Rambaut. “Genesis and pathogenesis of the 1918 pandemic H1N1 influenza A virus.” Proceedings of the National Academy of Sciences 111, no. 22 (2014): 8107–8112.

3. Nelson, Martha I., Cecile Viboud, Lone Simonsen, Ryan T. Bennett, Sara B. Griesemer, Kirsten St. George, Jill Taylor et al. “Multiple reassortment events in the evolutionary history of H1N1 influenza A virus since 1918.” PLoS pathogens 4, no. 2 (2008): e1000012

4. Lemon, Stanley M., and Adel AF Mahmoud. “The threat of pandemic influenza: are we ready?.” Biosecurity and bioterrorism: biodefense strategy, practice, and science 3, no. 1 (2005): 70–73.

5. Carter, Donald M., Christopher A. Darby, Bradford C. Lefoley, Corey J. Crevar, Timothy Alefantis, Raymond Oomen, Stephen F. Anderson et al. “Design and characterization of a computationally optimized broadly reactive hemagglutinin vaccine for H1N1 influenza viruses.” Journal of virology 90, no. 9 (2016): 4720–4734.

6. Dejnirattisai, Wanwisa, Daming Zhou, Helen M. Ginn, Helen ME Duyvesteyn, Piyada Supasa, James Brett Case, Yuguang Zhao et al. “The antigenic anatomy of SARS-CoV-2 receptor binding domain.” Cell 184, no. 8 (2021): 2183–2200.

7. Jackson, Cody B., Michael Farzan, Bing Chen, and Hyeryun Choe. “Mechanisms of SARS-CoV-2 entry into cells.” Nature reviews Molecular cell biology 23, no. 1 (2022): 3–20.

8. Yang, Chao-Fu, Chun-Che Liao, Hung-Wei Hsu, Jian-Jong Liang, Chih-Shin Chang, Hui-Ying Ko, Rue-Hsin Chang et al. “Human ACE2 protein is a molecular switch controlling the mode of SARS-CoV-2 transmission.” Journal of Biomedical Science 30, no. 1 (2023): 87.

9. Eswar, Narayanan, David Eramian, Ben Webb, Min-Yi Shen, and Andrej Sali. “Protein structure modeling with MODELLER.” Structural proteomics: high-throughput methods (2008): 145–159.

10. Teruel, Natália, Vinicius Magalhães Borges, and Rafael Najmanovich. “Surfaces: a software to quantify and visualize interactions within and between proteins and ligands.” Bioinformatics 39, no. 10 (2023): btad608.

11. Van Der Spoel, David, Erik Lindahl, Berk Hess, Gerrit Groenhof, Alan E. Mark, and Herman JC Berendsen. “GROMACS: fast, flexible, and free.” Journal of computational chemistry 26, no. 16 (2005): 1701–1718.

12. Case, David A., Hasan Metin Aktulga, Kellon Belfon, David S. Cerutti, G. Andrés Cisneros, Vinícius Wilian D. Cruzeiro, Negin Forouzesh et al. “AmberTools.” Journal of chemical information and modeling 63, no. 20 (2023): 6183–6191.

13. van Gunsteren, Wilfred F., and Herman JC Berendsen. “Algorithms for macromolecular dynamics and constraint dynamics.” Molecular Physics 34, no. 5 (1977): 1311–1327.

14. Valdés-Tresanco, Mario S., Mario E. Valdés-Tresanco, Pedro A. Valiente, and Ernesto Moreno. “gmx_MMPBSA: a new tool to perform end-state free energy calculations with GROMACS.” Journal of chemical theory and computation 17, no. 10 (2021): 6281–6291.

15. Ge, Jiwan, Ruoke Wang, Bin Ju, Qi Zhang, Jing Sun, Peng Chen, Senyan Zhang et al. “Antibody neutralization of SARS-CoV-2 through ACE2 receptor mimicry.” Nature communications 12, no. 1 (2021): 250.

16. Sievers, Fabian, and Desmond G. Higgins. “Clustal Omega for making accurate alignments of many protein sequences.” Protein Science 27, no. 1 (2018): 135–145.

