## Supplementary material for "How do we respond to the next SARS CoV epidemic/pandemic? A bioinformatics approach with the promise of preventing or reducing the severity of future SARS CoV related pandemics": Images and Tables

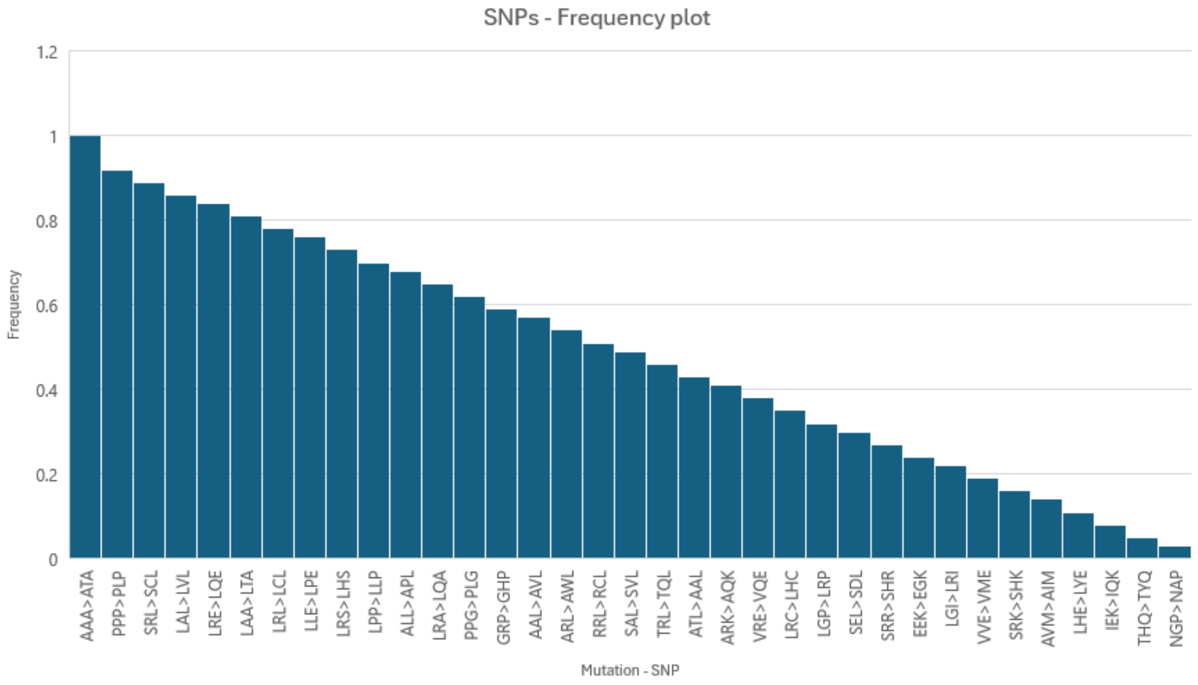


Fig.1 - Frequency plot of SNPs

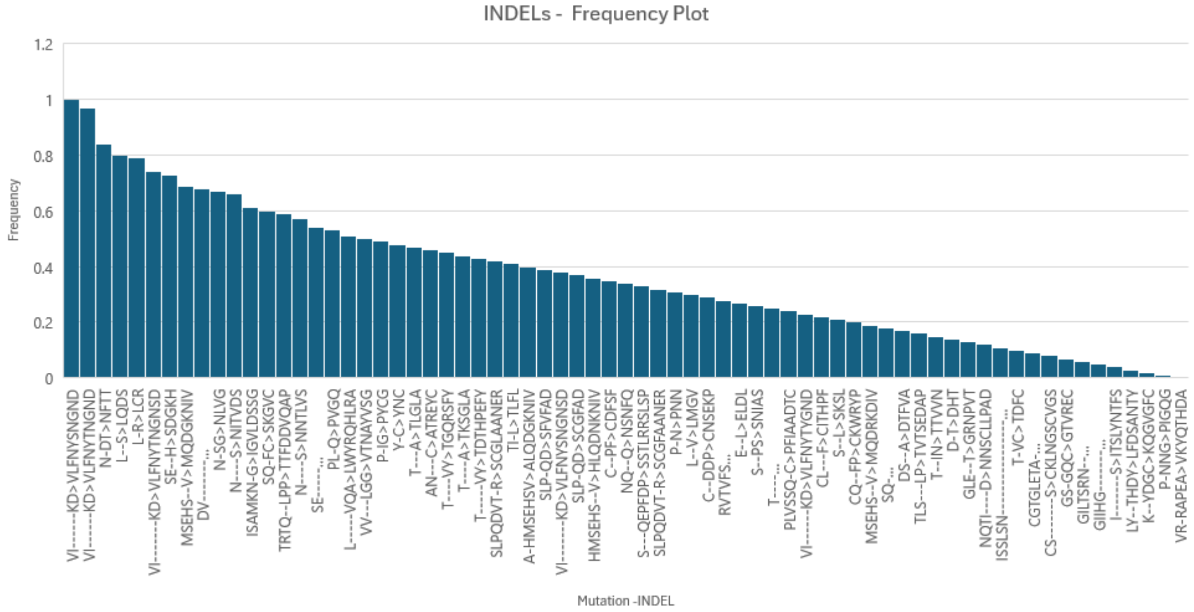


Fig.2 - Frequency plot of INDELs

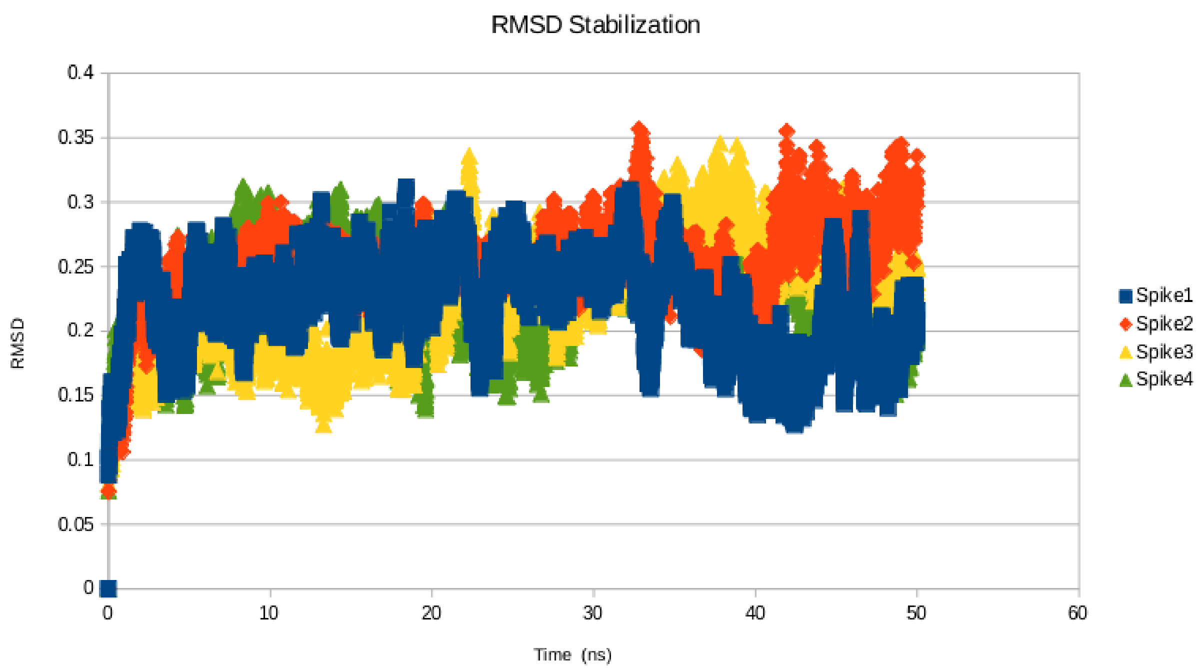


Fig 3 – RMSD stabilization indicating the stability of the structure corresponding to the chimeric Spike sequences generated through the approach.

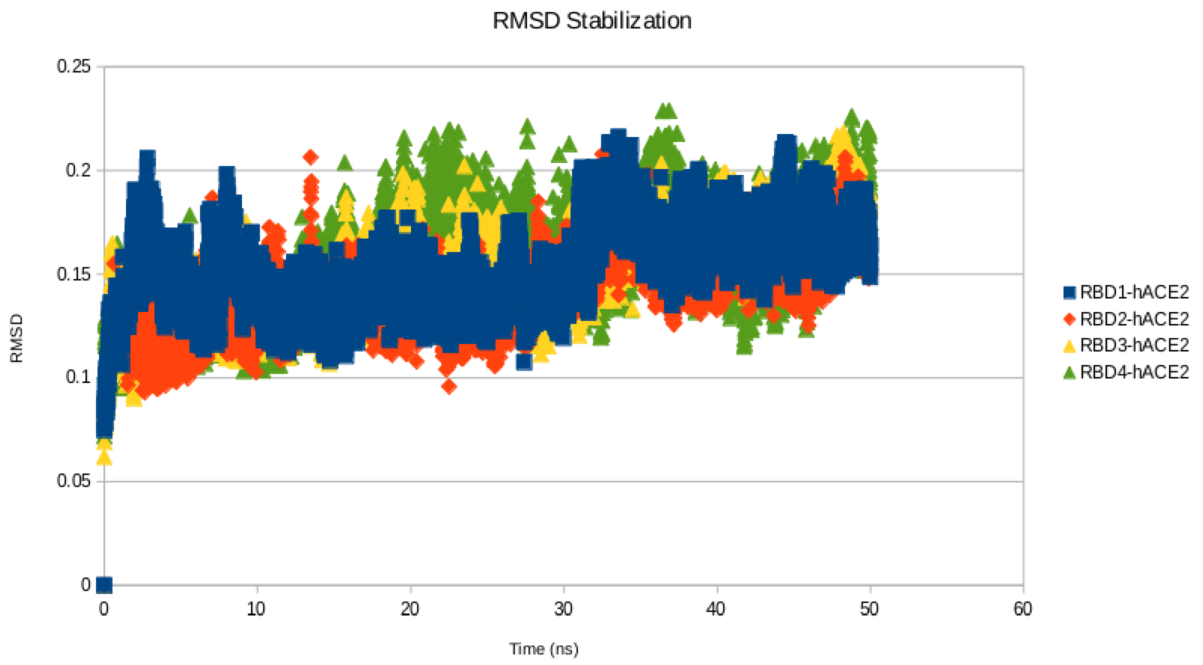


Fig.4 RMSD stabilization of RBD-hACE2 complexes corresponding to the Spike sequences generated by the Bayesian walker and shortlisted through the developed approach

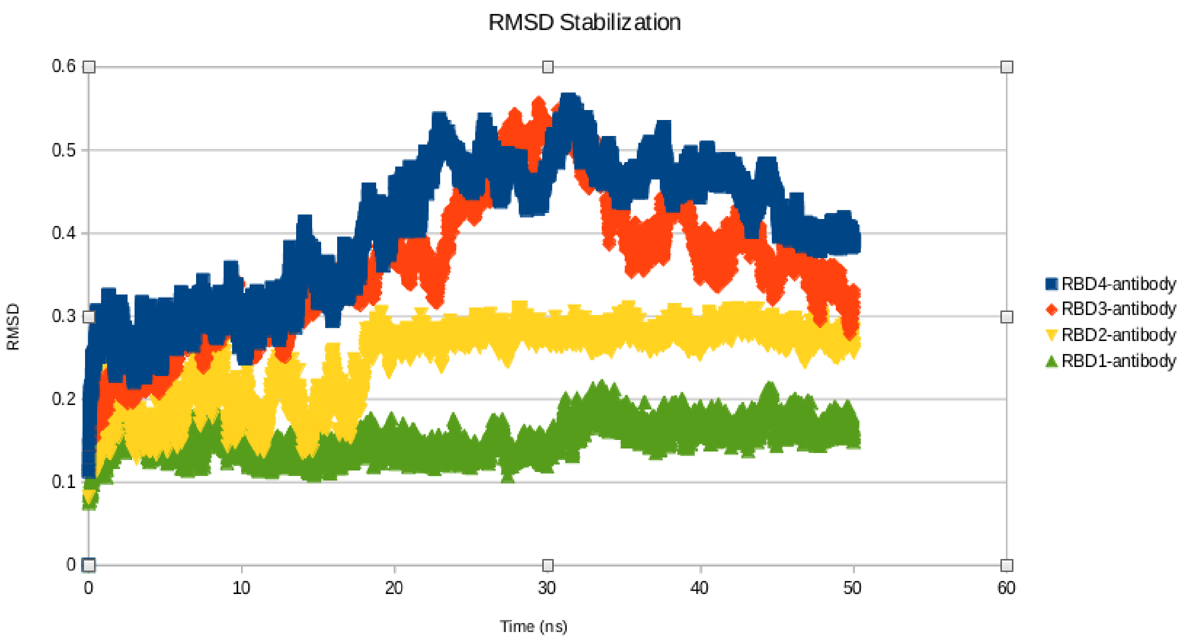


Fig.5 RMSD stabilization of RBD-antibody complexes corresponding to the Spike sequences generated by the Bayesian walker and shortlisted through the developed approach

```
SARS_CoV1_RBD       TNLCPFGEVFNATKFPSVYAWERKKISNCVADYSVLYNSTFFSTFKCY--
RBD1                -NLCPFGEVFNATKFPSVYAWERKKISNCVADYSVLYNSTFFSTFKCY--
RBD2                ---------------PSVYAWERKKISNCVADYSVLYNSTFFSTFKCY--
RBD3                ----------NATKFPSVYAWERKKISNCVADYSVLYNSTFFSTFKCY--
RBD4                ---------------------ERKKISNCVADYSVLYNSTFFSTFKCYRM
SARS_CoV2_RBD       TNLCPFGEVFNATRFASVYAWNRKRISNCVADYSVLYNSASFSTFKCY--
                                         :**:**************: *******

SARS_CoV1_RBD       --GVSATKLNDLCFSNV---YADSFVVKGDDVRQIAPGQTGVIADYNYKL
RBD1                --GVSATKLNDLCFSNV---YADSFVVKG-DVRQIAPGQTGVIADYNYKL
RBD2                --GVSATKLNDLCFSNV---YADSFVVKGDDVRQIAPGQTGVIADYNYKL
RBD3                --GVSATKLNDLCFSNVPLYYADSFVVKGDDVRQIAPGQTGVIADYNYKL
RBD4                GVGVSATKLNDLCFSNV---YADSFVVKGDDVRQIA-GQTGVIADYNYKL
SARS_CoV2_RBD       --GVSPTKLNDLCFTNV---YADSFVIRGDEVRQIAPGQTGKIADYNYKL
                      ***.********:**   ******::* :***** **** ********

SARS_CoV1_RBD       PDDFMG--CVLAWNTRNIDATSTGNYNYKYRYLRHGKLRPFERDISNVPF
RBD1                PDDFMG--CVLAWNTRNIDATSTGNYNYKYRYLRHGKLRPFERDISNVPF
RBD2                PDDFMGRVCVLAWNTRNIDATSTGNYNYKYRYLRHGKLRPFERDISNVPF
RBD3                PDDFMG--CVLAWNTRNIDATSTGNYNYKYRYL-HGKLRPFERDISNVPF
RBD4                PDDFMG---CLAWNTRNIDQG-----QYKYRYL-HGKLRPFERDISNVPF
SARS_CoV2_RBD       PDDFTG--CVIAWNSNNLDSKVGGNYNYLYRLFRKSNLKPFERDISTEIY
                    **** *    :***:.*:*       :* ** : :.:*:*******.  :

SARS_CoV1_RBD       SPDGKPCT-PPALNCYWPLNDYGFYTTT---GIGYQPYRVVVLSFELLNA
RBD1                SPDGKPCT-PPALNCYWPLNDYGFYTTT---GIGYQPYRVVVLSFELLNA
RBD2                SPDGKPCT-PPALNCYWPLNDYGFYTTT---GIGYQPYRVVVLSFELLNA
RBD3                SPDGKPCT-PPALNCYWPLNDYGFYTSTT-TGIGYQPYRVVVLSFELLNA
RBD4                SPDGKPCT-PPALNCYWPLNDYGFYYTTTGTGIGYQPYRVVVLSFELLNA
SARS_CoV2_RBD       QAGSTPCNGVEGFNCYFPLQSYGFQPTN---GVGYQPYRVVVLSFELLHA
                    .....**.   .:***:**:.***  :.   *:***************:*

SARS_CoV1_RBD      PATVCGPKLSTDLIKNQCVNF----------
RBD1                PATVCGPKLS---------------------
RBD2                PATVCGPKLSTDLIKNQCVNF----------
RBD3                PATVCGPKLSTDLI-----------------
RBD4                PATVC-PKLSTDLIKNQCVNFNFNGLTGTGV
SARS_CoV2_RBD       PATVCGPKKSTNLVKNKCVNF----------
                    ***** ** *
```

Fig.6 Alignment of RBDs

Table 1 Interaction and Binding energies of RBD-hACE2 complexes

| | Interaction score from Surface | Binding energy from gmx_MMPBSA Kcal/mol |
|---|---|---|
| RBD(SARS-CoV2)-hACE2 | -24.7 | -81.2 |
| RBD1-hACE2 | -23.73 | -80.7 |
| RBD2-hACE2 | -24.1 | -81.7 |
| RBD3-hACE2 | -24.3 | -81.5 |
| RBD4-hACE2 | -23.9 | -81.9 |

Table 2 Interaction and Binding energies of RBD-Antibody(P2C-1F11) complexes

| | Interaction score from Surface | Binding energy from gmx_MMPBSA |
|---|---|---|
| RBD(SARS-CoV2)-Antibody(P2C-1F11) | -24.9 | -83.7 |
| RBD1- Antibody(P2C-1F11) | -11.9 | -22.7 |
| RBD2- Antibody(P2C-1F11) | -7.6 | -15.5 |
| RBD3- Antibody(P2C-1F11) | -5.7 | -13.7 |
| RBD4- Antibody(P2C-1F11) | -5.9 | -11.1 |
